# Rotational snapping: Illusory rhythmicity induced by global and local motion binding

**DOI:** 10.1101/2020.02.03.931758

**Authors:** Hio-Been Han

## Abstract

The complexity of human perceptual system has often been investigated through its atypical function; the illusion. Here, I introduce a new visual illusion induced by rotational motion binding, which leads to a gestalt perception of illusory object rhythmically popping out and fading away. In this illusion, observers reported non-existing rhythm out of rotating Gabor patches (*e.g*., local rotation) that also have orbital trajectory with opposite direction (*e.g*., global rotation), only under the particular combinations of parameters. This illusory rhythmicity was four times faster than the average of global/local rotational speed. Image reconstruction using the response-triggered average revealed the rhythmicity is explained by the circular alignment of array, demonstrating the effect of repeated contour integration and its perceptual consequences.

## INTRODUCTION

When clockwise-rotating array composed of counter-clockwise-rotating Gabor patches at certain speed, perceived rhythm approximately four times faster is induced, if the patches have particular phase disparity of rotation (Fig 1A, Supplementary Movie 1A) This rhythmicity is an illusion because the rhythmicity doesn’t exist in any component of the rotation, hinting at the dynamic binding process in visual system. Through a pilot experiment (n=7), I found the various components of the rotational motion were all crucial for the effect. Schematic illustrations of four control stimuli (*‘No global motion’, ‘No local rotation’, ‘+1/2π phase lag’, ‘Same directional rotation*) are shown in Fig 1B whose four-time-faster rhythmicity were reported from none of the observers (See Supplementary Movie 1B-E). To figure out why do we perceive the illusory rhythmicity from it, a psychophysical experiment was conducted.

**Figure 1.**
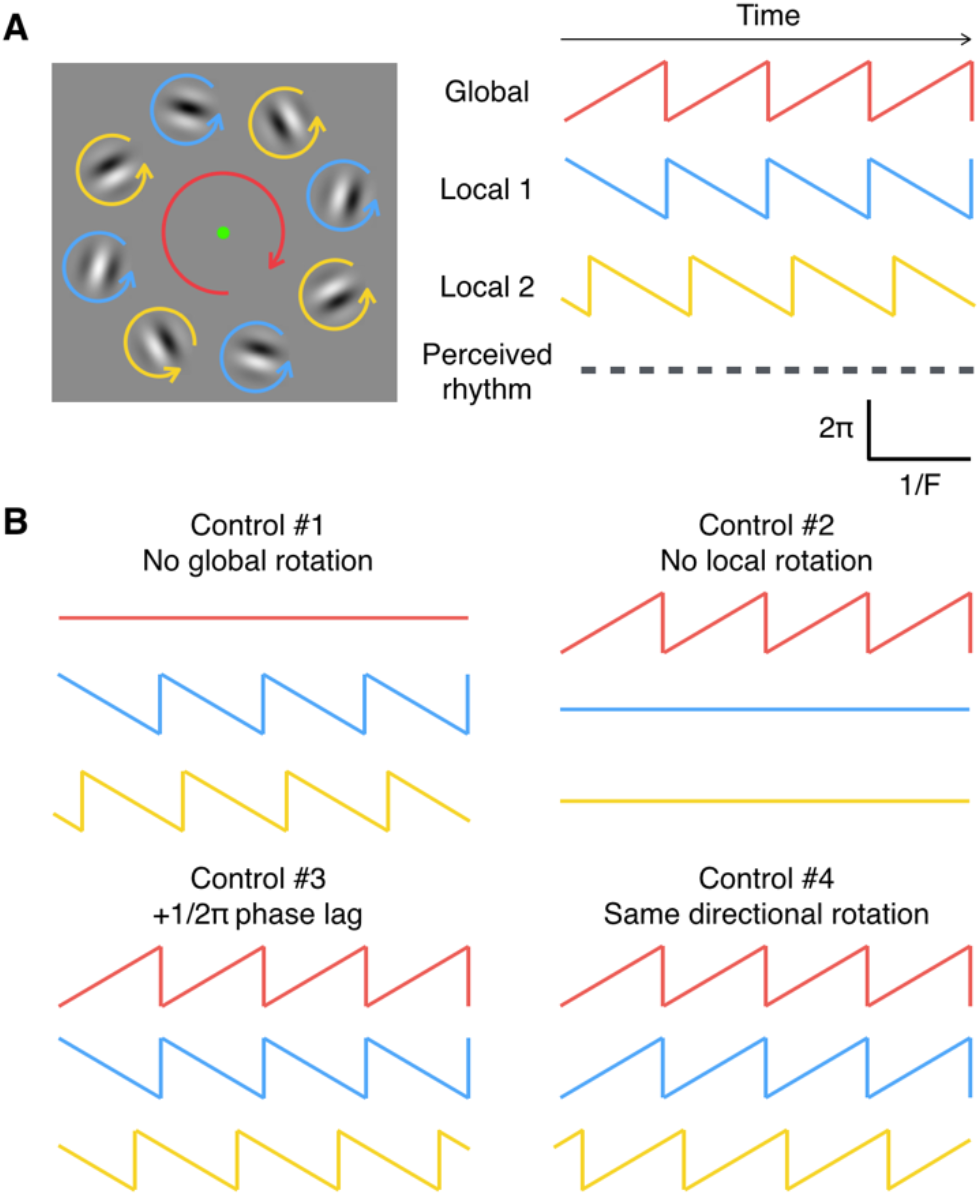
Schematics of rotational motion components. (A) Display condition which induced illusory rhythm. (B) Display conditions which didn’t induced the rhythm.

## RESULTS

Recruited observers (n=3) were seated in front of computer monitor then asked to press keyboard button repeatedly when they find rhythmic change or transitions from the video frames. In the video frames, eight Gabor stimuli belonging either to one of two subgroups (phase lag between groups = −1/2π, Fig 1A) rotating at various speed (five local-rotation conditions; 0.40, 0.45, 0.50, 0.55, 0.60 Hz, counterclockwise), while the array was rotating (five global-rotation conditions; 0.40, 0.45, 0.50, 0.55, 0.60 Hz, clockwise). The sample video frames are summarized in Supplementary Movie 2. During the presentation of 12 sec movie frames (total 75 trials; 3 trials * 25 conditions), observer’s binary responses were sampled (Fig 2A) at frame rate (60 Hz) and analyzed in time-frequency domain (Fig 2B-C).

**Figure 2.**
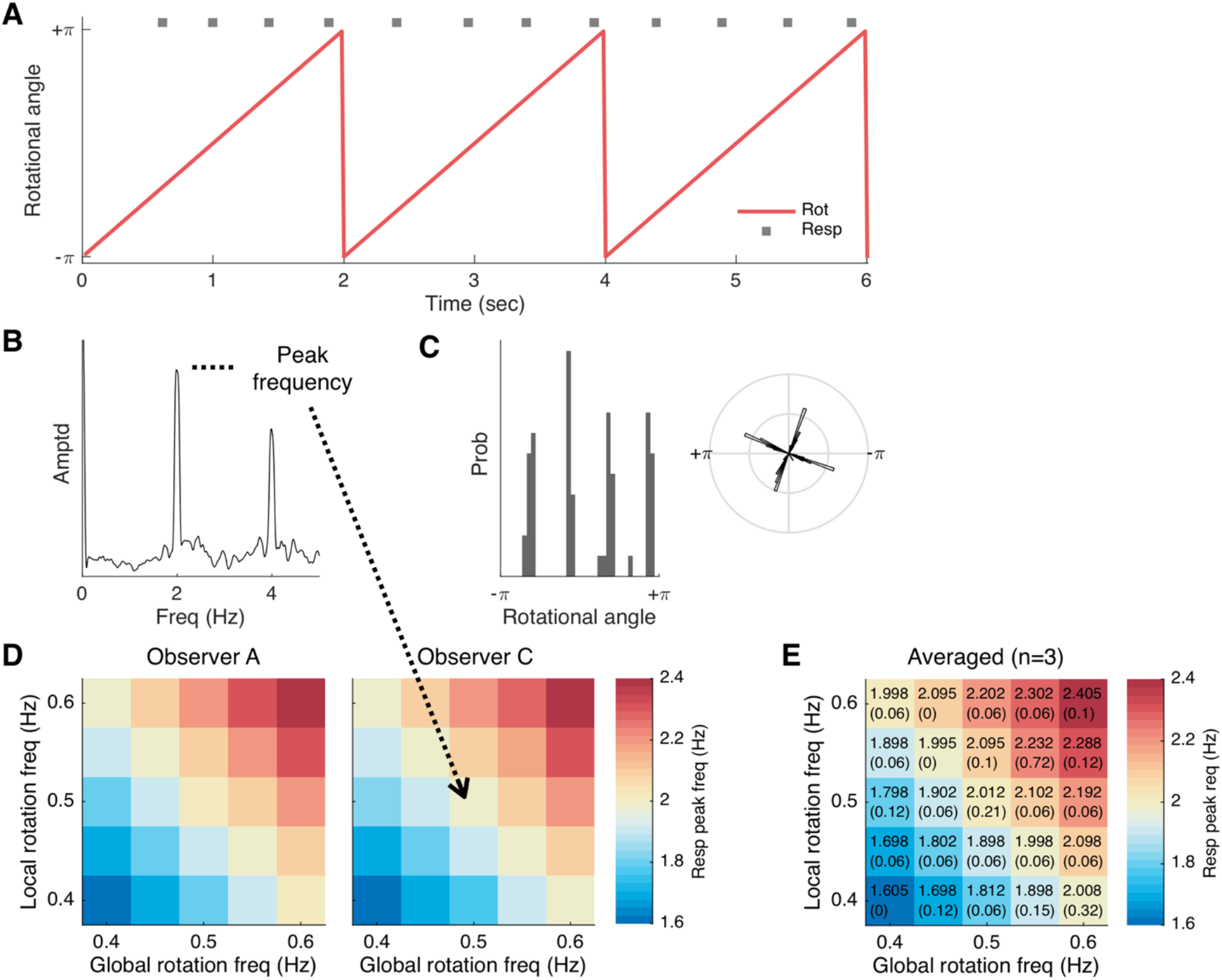
Rotational motion binding and induced rhythmicity. (A) Example time trace of motion (global/local motion at 0.5/0.5 Hz, respectively) and key responses (near 2 Hz). (B) Absolute amplitude of frequency spectrum of key response. (C) Response histogram relative to rotational angle. (D, E) Peak frequency detected in given response from observer(s). Values in each pixel denote the mean (standard deviation) peak frequencies in (E).

### Illusion speed determined by rotation speed

To reveal the temporal relationship between rotation and perceived rhythmicity, peak frequency in amplitude spectral density (*i.e*., 2 Hz in case of Fig 2B) was obtained. Fig 2D and 2E illustrate the measured illusion frequency as a function of rotational frequency. When the global and local rotation have the same speed (*i.e*., global 0.5 Hz, local 0.5 Hz), illusion frequency showed four times faster than the speed (*i.e*., 2.0 Hz). Interestingly, when they have different speed (*i.e*., global 0.5 Hz, global 0.6 Hz), illusion frequency was equal to four times faster than their average speed (*i.e*., 2.2 Hz). This relationship was violated in none of the global/local frequency combination, all *P*s >.25 (uncorrected Wilcoxon signed-rank tests, the null hypothesis was set to that response frequency is equal to mean rotational frequency times four).

What induced the rhythm? One possibility might be that spatial alignment of stimuli forms a contour-integrated object (*i.e*., circle, square, etc.) periodically. Confirming this hypothesis, I performed simple reverse correlation by taking response-triggered averaging (RTA) image from video frames to capture the characteristics of stimuli’s spatial alignments as a function of response frequency.

### Regular alignments of circular contours

Individual RTA image at each time point was obtained. For this, time bin for sampling were rescaled across the conditions because illusion speed varies, and then set to 15-step bins covering 2.5 cycles (−2π to +3.5π) of response frequency (*i.e*., from −500 to 875 ms for 2 Hz responses, from −555.56 to 972.22 ms for 1.8 Hz responses, etc.). As RTA images from odd/even responses showed inversed contrast (see Fig 3A, mid: odd, top & bottom: even), they analyzed separately. First and few more responses were discarded to set the number of responses be the multiple of 4, preventing odd/even orders being mixed up. After obtaining RTA image of each condition, image contrast was calculated using root-mean-square (RMS) contrast (Peli, 1990) then transformed into z-score to control the difference of dynamic range across the conditions.

**Figure 3.**
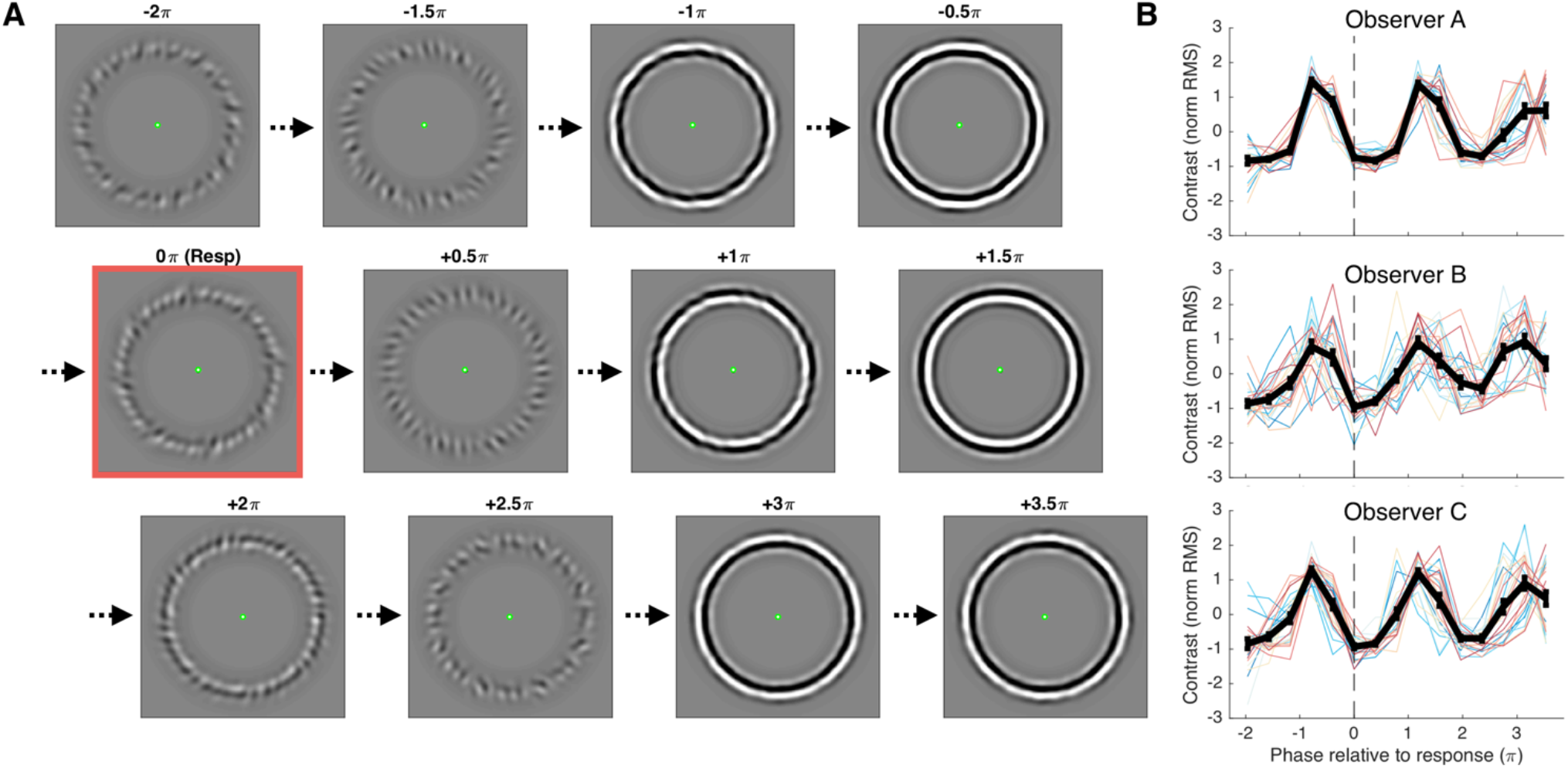
Regularity of array alignments captured by reverse correlational response-triggered average. (A) Grand-averaged RTA images of observer A across all trials (n=75) relative to illusion frequency (4X of mean rotation speed) from odd key presses (n=662). Red square highlights RTA image from moment of the response (Top row: N-1th cycle, mid row: Nth cycle, bottom row: N+1th cycle). (B) Periodic fluctuation of RTA image contrast measured from 3 observers. Each colored solid line indicates 1 out of 25 global/local speed combinations, black dashed line indicates moment of the response (odd), and black solid line indicates the mean contrast value across the conditions as a function of relative phase to illusion frequency. Error bars represent standard error of the means.

As the Gabor patches are composed of black, gray and white values with an orientation, RTA image should have low contrast unless patches are aligned with a bias toward a particular orientation across responses/trials. Thus, the sharper RTA image implies the higher probability of arrays’ alignment at given phase on response cycles. To check the alignment regularity, the contrast was averaged in each time point (*i.e*., point of phase relative to response frequency) (solid black lines in Fig 3B). I could find a clear tendency of rhythmic fluctuation that contrast in RTA images changes over time, as shown in Fig 3B. Simple curve-fitting with one-term sinusoidal function with illusion frequency explained major portion of variations from the data (odd response *R^2^* = .85, .83, .95; even response *R^2^* = .87, .87, .95; for observer A, B, C, respectively). In other word, arrays’ alignment—forming a closed circle shown in Fig 3A—was quite regular at some particular phase of response cycle. Combining with the closed-contour circular shape observed in RTA image, this result suggest the circle composed of Gabor-contours might be the hidden substance of rhythmicity.

## DISCUSSION

In this article, a novel illusion was introduced that rotational motion binding produced illusory rhythmicity. I showed that the speed of rhythm was completely determined by the mixture of global and local rotations (Fig 2), and that the rhythm can be explained by a consequence of periodic contour integration process forming circular-shape closed object (Fig 3).

One interesting aspect of this illusion is that it’s a consequence of actual *binding* process (*i.e*., multiple motion signals). For instance, the similar effect was not induced without global rotation (Supplementary Movie 1B) even though the moments of circular alignment still exist. This suggests the illusory rhythm reflects dynamic and complex characteristic of motion processing in the human visual system.

### Relevance to perceptual closure effect

Feature integration is one of the most fundamental processes in human visual cognition (Treisman, 1988). The illusory rhythm appeared here might be explained in a way of gestalt principles such as perceptual closure or grouping which follows “good-continuation” law of gestalt psychology, leading to transient popping-out of the salient circular object (Wertheimer, 1938; Barlett, 1916). In this point of view, the illusion introduced here may reflect temporal aspect of “association field” proposed by modern gestalt psychologists (Field *et al*., 1992; Hess *et al*., 2003).

### The 4X speed of illusory rhythmicity

Another interesting part of this illusion is that the rhythm observers *psychologically* perceived actually doesn’t *physically* exist in the single rotations. Why the rhythm gets the 4X speed? One explanation could be the frequency integration. In the visual stimuli, physically-existing frequencies were two: *f*_1_, *f*_2_ (for global, local, respectively). When they are summed up, it makes a new frequency, *f*_1_ + *f*_2_. However, the observers reported doubled speed of it, 2(*f*_1_ + *f*_2_), not the summed frequency *per se*.

What the observers were looking at, in fact, was two different streams with speed of *f*_1_ + *f*_2_. The only reason they perceived doubled speed is that those streams have anti-phasic relationship. As Fig 3A shows, separating RTA images from odd/even responses, whose color got reversed (*i.e*., inner-black outer-white circle and inner-white outer-black circle), makes two *f*_1_ + *f*_2_ streams with half-cycle-shifted phase relationship. To confirm this idea, simplified control stimuli were made and it shows the two *f*_1_ + *f*_2_ streams speed separately (See Supplementary Movie 3). As expected, this separation of image streams induced two anti-phasic distinctive rhythms speed of 2X (*i.e*., 1 Hz rhythm from 0.5 Hz rotations). In this regard, the 4X-speed rhythmicity observed in the experiment can also be explained by simple superposition of the waves (*i.e*., traveling wave).

## Conclusion

The illusion introduced here is related to not only the contour integration but also the dynamic motion binding process in the visual system. Future studies may adopt this illusion to explore the mysterious nature of contour integration as well as motion binding processing. Furthermore, the illusion introduced here may be useful tool for perceptual processes in rodent model as it doesn’t require verbal nor behavioral report, replacing typical drifting-grating or random-dot stimuli (Han *et al*., 2017; Han *et al*., 2019).

## METHODS

Three observers (67% female, age *M* = 23.33, *SD* = 2.08) with normal vision participated the experiment. None of the procedure was approved by the Institutional Review Board of Korea Advanced Institute of Science and Technology (KAIST).

### Materials

Visual stimuli were generated with MATLAB R2015b (Mathworks, MA, USA) and presented via Psychophysics Toolbox (Brainard, 1997) v3.0.14 on MacBook Pro Retina 13.3’ monitor at 60 Hz. Observers viewed the stimuli from 41 cm apart from the monitor (Gabor spatial frequency = 1.4 cycle/°, Gabor radius = 2.1°, global field radius = 11.6°), and asked to response by pressing left shift key on the keyboard. Each trial consisted of 12 s of video frames and following 2-3 s of inter trial interval. Green dot central fixation (radius = 0.15°) was always presented.

### Data analysis and statistical tests

Binary pulse train of key press data were transformed into frequency domain using fast Fourier transform, then interpolated and smoothed to detect the peak response frequency more accurately. Non-parametric t-test was performed using Statistic and Machine Learning Toolbox v10.1 of MATLAB to test whether peak frequency is different from average rotational frequency times four or not (α = 0.05). Fluctuation of contrast value from RTA images over instantaneous phase of response frequency was fitted on single-term sinusoidal function using Curve Fitting Toolbox v3.5.2 in MATLAB.

## Supporting information

Cover Video

## SUPPLEMENTAL INFORMATION

Supplementary movies are available online:

Supplementary Movie 1. https://youtu.be/egyIcpPG03o

Supplementary Movie 2. https://youtu.be/iGlbcrq4j0c

Supplementary Movie 3. https://youtu.be/-JjMEtHHreU

## ACKNOWLEDGEMENTS

I specially thank to Prof. Michael Bach for his naming suggestion of this illusion.

